# Adapting to novel environments together: evolutionary and ecological correlates of the bacterial microbiome of the world’s largest cavefish diversification

**DOI:** 10.1101/2021.11.18.469109

**Authors:** Shipeng Zhou, Amrapali Prithvisingh Rajput, Yewei Liu, Tingru Mao, Jian Yang, Jayampathi Herath, Madhava Meegaskumbura

## Abstract

The symbiosis between a host and its microbiome is essential for host fitness, and this association is a consequence of the host’s physiology and habitat. *Sinocyclocheilus*, the largest cavefish diversification of the world, an emerging multi-species model system for evolutionary novelty, provides an excellent opportunity for examining correlates of host evolutionary history, habitat, and gut-microbial community diversity. From the diversification-scale patterns of habitat occupation, major phylogenetic clades (A–D), geographic distribution, and knowledge from captive-maintained *Sinocyclocheilus* populations, we hypothesize habitat to be the major determinant of microbiome diversity, with phylogeny playing a lesser role. For this, we subject environmental water samples and fecal samples (representative of gut-microbiome) from 24 *Sinocyclocheilus* species, both from the wild and after being in captivity for six months, to bacterial 16S rRNA gene profiling using Illumina sequencing. We see significant differences in the gut microbiota structure of *Sinocyclocheilus,* reflective of the three habitat types; gut microbiomes too, were influenced by host-related factors. There is no significant association between the gut microbiomes and host phylogeny. However, there is some microbiome related structure at clade level, with the most geographically distant clades (A and D) being the most distinct, and two geographically overlapping clades (B and C) being similar. Microbes inhabiting water were not a cause for significant differences in fish-gut microbiota, but water quality parameters was. Transferring from wild to captivity, the fish microbiomes changed significantly and became homogenized, signifying adaptability and highlighting the importance of environmental factors (habitat) in microbiome community assembly. The core microbiome of this group closely resembled that of other teleost fishes. Our results suggest that divergent selection giving rise to evolutionary novelties also includes the microbiome of these fishes, which provides a functional advantage for life in the resource-depleted cave environment.

**SIGNIFICANCE STATEMENT:** The largest diversification of cavefishes of the world, *Sinocyclocheilus*, not only show that habitat, and phylogenetic clade is important in determining their gut microbiome, but also that they reach a common microbiome in captivity irrespective of their phylogenetic position, region of origin and habitat, indicating that they are adaptable in the context of microbe related changes in their environment.

## INTRODUCTION

The gastrointestinal tract of an animal is occupied by a microbiome, a staggering diversity of microbial colonies that are in a symbiotic association with the host (Hooper, 2015). These commensal gut-bacterial relationships influence many vital aspects of the host, such as immune function, nutrient absorption, development, and behavior (Mazmanian *et al.*, 2005; Nicholson *et al.*, 2005; Mcfall-Ngai *et al.*, 2013). Hence, the gut microbiome is considered an extension of the host genome, and these microbes are thought to have served as a “bridge” between the host and the external environment during evolution (Shapira and Michael., 2016). Therefore, the microbiome of an organism, an adaptation contributing to fitness, is the result of an interplay between the host’s phylogeny and its environment (Gill *et al.*, 2006). In large diversifications, where hosts derive from a common ancestor to form clades and diversify to occupy distinct habitats and regions, some of these ideas related to microbiome community assembly can be tested.

The phylogenetic relationships of an organism has an influence on the structure of its microbiome (Brucker and Bordenstein, 2012). The term “phylosymbiosis” has been used to encapsulate this idea, which has been supported through controlled laboratory experimentation and analyses of wild populations (Brooks *et al.*, 2016; Kohl *et al.*, 2017; Ingala *et al.*, 2018). However, large diversification-scale studies addressing correlates of host evolutionary history and microbial community diversity are still lacking.

A related idea extending from phylosymbiosis is that of the “core-microbiome”, which was initially introduced in a narrower sense in human host studies (Turnbaugh *et al.*, 2009; Qin *et al.*, 2010; Turnbaugh *et al.*, 2010). Since then, a plethora of core-microbiomes, ranging from single species to higher taxonomic groups, has manifested, reinforcing the association between host phylogeny and gut microbiome composition. The selective-mechanisms that contribute to the establishment of a core-microbiome in hosts, however, are unclear. Differential selection pressures in the form of host-physiology (linked to host phylogeny) and environment on microbiota in large diversifications occupying contrastingly different habitats, can shed light on gut-microbe and host associations. Such a system exists in species that have evolved to occupy caves.

Various vertebrate groups have occupied caves by adapting to conditions of low availability of food, oxygen and light, leading to repeated evolution of troglomorphic adaptations such as lowered metabolism, specialized behaviors, specialized sensory systems, and loss of eyes and pigmentation (Zhao and Zhang, 2006; Gross *et al.*, 2008). This accelerated enhancement or regression of certain traits in troglomorphic forms is a useful model system to study the process of natural selection in response to cave-associated selective regimes (Yang *et al.*, 2016; Hart *et al.*, 2020; Zhao *et al.*, 2021). Though rarely tested, cave systems provide opportunities to test hypotheses pertaining to microbiome community assembly in response phylogeny and environment. Although large troglomorphic diversifications are rare, one such radiation exists in a group of freshwater fishes in China. *Sinocyclocheilus* (Cyprinidae, Barbinae), the largest cavefish diversification in the world, represents an emerging model system for evolutionary novelty (Mao *et al.*, 2021), provides an invaluable opportunity for examining the associations between host evolutionary history, environment and gut-microbial community diversity. With almost 75 extant species, *Sinocyclocheilus* is a monophyletic group of cyprinid cavefishes endemic to the expansive southwestern karstic region of China, the largest limestone area in the world, with an area ca. 620,000 km^2^ (Jiang *et al.*, 2019). Having first evolved during the late-Miocene, these fishes show morphology-to-habitat correlation in a staggering array of adaptations to subterranean life. Interestingly, they also show several independent events of cave occupation. The microbiome of *Sinocyclocheilus* has not been subjected to a diversification-scale study so far.

Recent mt-DNA based phylogenies of *Sinocyclocheilus* suggest four major clades, Clades A – D, each with distinct morphologies, habitat-occupation strategies and geographic ranges. They extend from the Eastern Guangxi autonomous region to the South-Eastern Guizhou and Eastern Yunnan Provinces of China. The earliest emerging clade (Clade A) is restricted to Guangxi, at the Eastern fringes of the distribution of the genus. Clades B and C, which have overlapping distributions, are restricted to the middle of the range of the genus (Guizhou, North-Central and North-Western Guangxi), and species of Clade D are found mostly in lotic habitats associated with hills to the west. They are classified into three major habitat types as Troglobitic (exclusively cave species), Troglophilic (associated with caves), and Surface (non-cave dependent species). These habitat types are correlated with their morphological adaptation, especially of their eye condition: Normal-Eyed (Surface), Micro-eyed (Troglophilic) and Blind (Troglobitic) (Mao *et al.*, 2021).

While earlier gastrointestinal tract related microbiome studies focused mostly on fishes of economic importance, recent studies have focused on species of evolutionary, ecological or conservation significance (Talwar *et al.*, 2018). *Sinocyclocheilus* is identified with the later (recent) group of studies. However, very little is known of the feeding ecology and the microbiome of *Sinocyclocheilus.* Cave-inhabiting species feed on cave insects, bat excrement and debris carried by the water flow (Zhang *et al.*, 2015). A previous study on the gut microbiota of *Sinocyclocheilus* showed abundance of cellulose-degrading bacteria such as *Bacillus*, *Clostridium*, and *Planctomyces,* in the cave dwelling species compared to the surface species (Chen *et al.*, 2019).

Water chemistry has been associated with the gut microbiome of hosts in a different cavefish system. The dissolved oxygen content in cave water was shown to alter the β-diversity of gut microbiota in *Astyanax mexicanus*, the well-known single-species cavefish model system (Ornelas-Garcia *et al.*, 2018). This suggests that cavefish species and populations have distinct microbial communities to facilitate life in the subterranean environment. However, such data do not exist for *Sinocyclocheilus.*

Furthermore, the changes of the bacterial microbiome when cavefish populations are transferred from various wild habitat types to captivity has not been documented so far. This can be used to study the changes in microbiome community in the short run, helping further understand the acquired microbiome for the life of these fish and the factors that may influence such microbiome assembly. Knowledge of the changes in the microbiome is also useful for conservation breeding, reintroduction and study of these rare and threatened cavefishes. As a part of our explorations into cavefishes, we have maintained 40 *Sinocyclocheilus* species in captivity from 1-3 years, with many of them still thriving (Pers. Obs.). Since the microbiome is dependent on external factors, such as water conditions and diet, we presume, given proper care, that they are adaptable, together with their gut microbiomes.

Here we carry out a diversification-scale analysis of *Sinocyclocheilus* using 24 representative species of the phylogeny (phylogeny has a strong association with geography) and habitat representation to understand the host-microbiome relationships of these fishes, including that in captivity. We formulate our hypothesis based mainly on the following observations: (□) these fishes have repeatedly evolved to occupy caves, independent of the phylogeny (□) given appropriate water conditions and diet, these fishes can survive in captivity over long periods, hence they and their microbiome are adaptable. From these observations, we hypothesize that the microbiome of *Sinocyclocheilus* is dependent primarily on habitat associations (Troglobitic – Blind; Troglophilic – Micro-Eyed; Surface – Normal), with phylogeny playing a secondary role. We test this hypothesis through the following predictions: (□) That the microbiomes of troglobitic species are more similar to one another than to surface species, with the troglophilic species in between. Since there are independently derived blind (troglobitic) species, we predict that they will have similar microbiomes. (□) That the microbiomes of the four clades will be similar, except clades B and C as they overlap in geographic distribution. (□) Since *Sinocyclocheilus* is monophyletic, that there will be a core microbiome characteristic of the group. (□) Since we have maintained these fishes in captivity for more than three years, we predict that they can survive on an acquired microbiome that is different from the wild one.

## RESULTS

We collected 24 species of *Sinocyclocheilus* from the wild, together with their fecal samples and water samples from their habitat, they were then raised in captivity (Figure 1). After six months in captivity, we sampled their feces again (17 species). Our sequence library contained microbiomes from 75 wild fecal samples, 46 captive fecal samples, and 19 water samples (Supplementary Table S1). According to the rarefaction curves, the estimations of species richness were steady and unbiased for all samples (Supplementary Figure S1).

**FIGURE 1.**
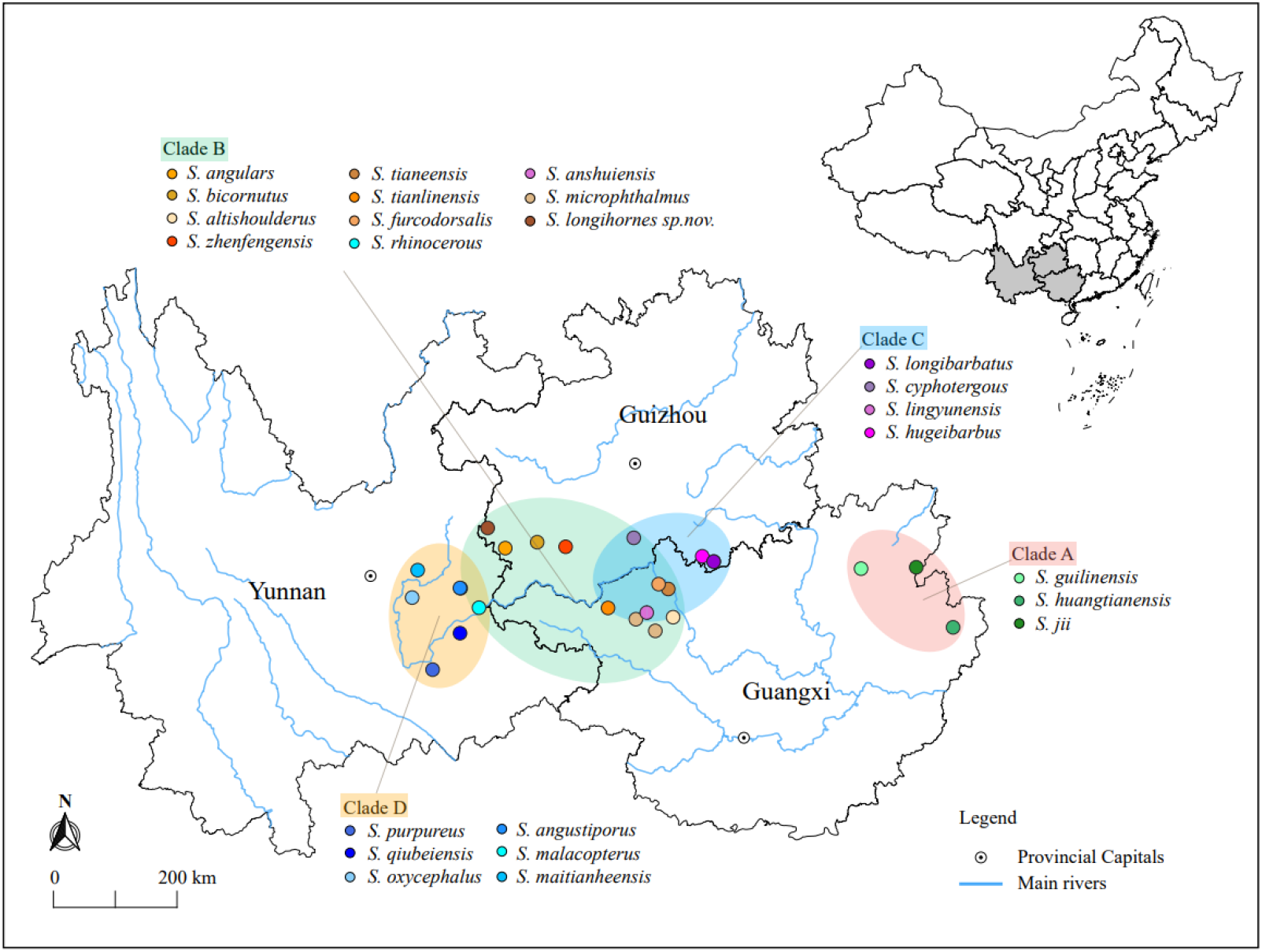
Distribution of 24 *Sinocyclocheilus* species and geographic context of phylogenetic clades. Colored dots represent the species. The four phylogenetic clades are represented by the colored ellipses. Clades A (East) and D (West) are disjunct and Clades B and C are overlapping.

### Composition of gut microbiomes in *Sinocyclocheilus*

We used wild populations to demonstrate the natural gut microbial community structure of cavefish (Supplementary Table S2). The predominant phyla composing the gut microbes of *Sinocyclocheilus* were Proteobacteria, Fusobacteria, Firmicutes, Bacteroidetes, and Verrucomicrobiota, with relative abundances of 57.4%, 31.6 %, 6.1%, 2.7%, and 0.4%, respectively. As expected, there are differences between *Sinocyclocheilus* species, mainly in the abundance of Proteobacteria and the Fusobacteria (Figure 2). Individual species in the same clade (Clade A, Clade C and Clade D) had similar gut microbiota, while individual species in Clade B had a diverse gut microbiota (Figure 2). At level of taxonomic order, considering all wild-samples, the most abundant taxa were Fusobacteriales (31.6%), Aeromonadales (32.7%), Enterobacterales (10.9%), Burkholderiales (5.4%), Pseudomonadales (2.9%) and Bacteroidales (1.8%) (Supplementary Figure S2a). In addition, the gut microbiota of all these cavefish were dominated by eight genera (Supplementary Figure S2b), *Cetobacterium* (31.6%), *Aeromonas* (31.2%), *Acinetobacter* (2.2%), *Shewanella* (1.5%), *ZOR0006* (1.4%), *Deefgea* (1.3%), *Enterobacter* (1.0%) and *Crenothrix* (0.9%).

**FIGURE 2.**
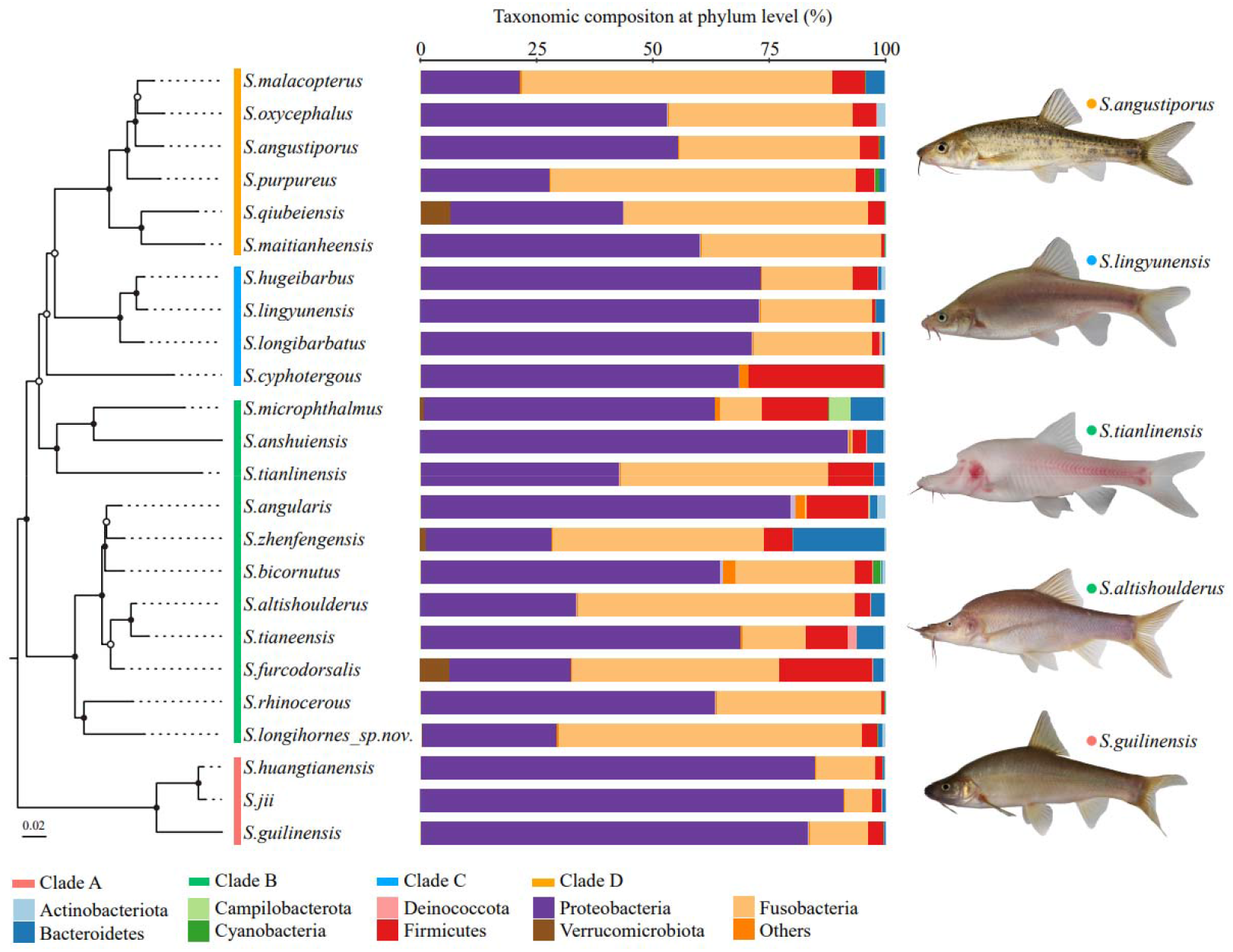
Phylogenetic relationships and microbial composition of *Sinocyclocheilus* species (N = 24). Left: The maximum likelihood phylogenetic tree based on concatenated *cytb* and *ND4* gene fragments of 24 *Sinocyclocheilus* species, with 1000 bootstrap replications. Filled circles on the nodes denote ≥ 70% bootstrap support; Middle: Phylum level composition of the gut microbiome. The microbial community for each species is displayed as the sum of all individuals within a given species; Right: Images of representative species in four phylogenetic clades.

### The association of gut bacterial communities with the host phylogeny

We used Mantel tests to assess whether between-species gut microbial distance is correlated with between-species genetic and geographic distances. There was no significant correlation between-species gut microbiota and geographical distance (Bray-Crutis Mantel *R* = −0.035, *P* = 0.568; unweighted UniFrac *R* = 0.065, *P* = 0.23; weighted *R* = 0.065, *P* = 0.263). And the gut microbiota and *Sinocyclocheilus* genetic divergence were insignificant irrespective of geographical distance (Bray-Crutis Mantel *R* = −0.119, *P* = 0.823; unweighted UniFrac *R* = −0.038, *P* = 0.601; weighted *R* = −0.195, *P* = 0.895). To test for signatures of host phylogeny on metagenomic community composition, we first constructed phylogenetic trees using the maximum likelihood estimation for *cytb* and *ND4* genes from 24 species of *Sinocyclocheilus*, and then used unweighted Unifrac distance matrices to construct UPGMA trees. We found that the phylogenetic tree and the similarity distance clustering tree appeared to be unrelated (Bray-Crutis, Robinson-Foulds distance = 36.0; unweighted UniFrac, Robinson-Foulds distance = 38.0; weighted UniFrac, Robinson-Foulds distance = 38.0).

Next, we investigated the variation of intestinal flora at a lower scale, i.e., under different phylogenetic clades and habitats. We classified wild populations of *Sinocyclocheilus* into four groups based on their clades and three groups based on their habitats, which have been reported by other authors (Yang *et al.*, 2016; Mao *et al.*, 2021) (Supplementary Table S1). Simpson index revealed that within-sample diversity (alpha diversity) of Clade D differs from those of Clade A and Clade B, while Clade C was not significantly dissimilar from the other groups (Figure 3a and Supplementary Table S3). Overall, the gut microbiota of Clade A and Clade B had higher diversity than that of Clade D. Furthermore, a Constrained Principal Coordinate Analysis (CPCoA) of the full dataset revealed that the host phylogeny has a significant impact on bacterial communities, accounting for 11.5 % of the variation (Figure 3b). We found a clear separation between Clade A and Clade D from the first component, as well as an overlap between Clade B and Clade C (Figure 3b). It indicated that Clade A and Clade D were each inhabited by a distinctive bacterial community, while the Clade B and Clade C had similar microbiomes. Further, the metagenomic relationships of four groups inferred by computing unweighted Unifrac distance matrices indicated that the microbial composition of the four clades were consistent with phylogenetic relationships (unweighted UniFrac, Robinson-Foulds distance = 0.0) (Figure 3c). Bacterial composition varied across clades at the phylum level, as evidenced by a decrease in Proteobacteria and an increase in Fusobacteria from A to D (Figure 3c). However, this change was not linear, as pattern for Clade C changed before Clade B (Figure 3c).

**FIGURE 3.**
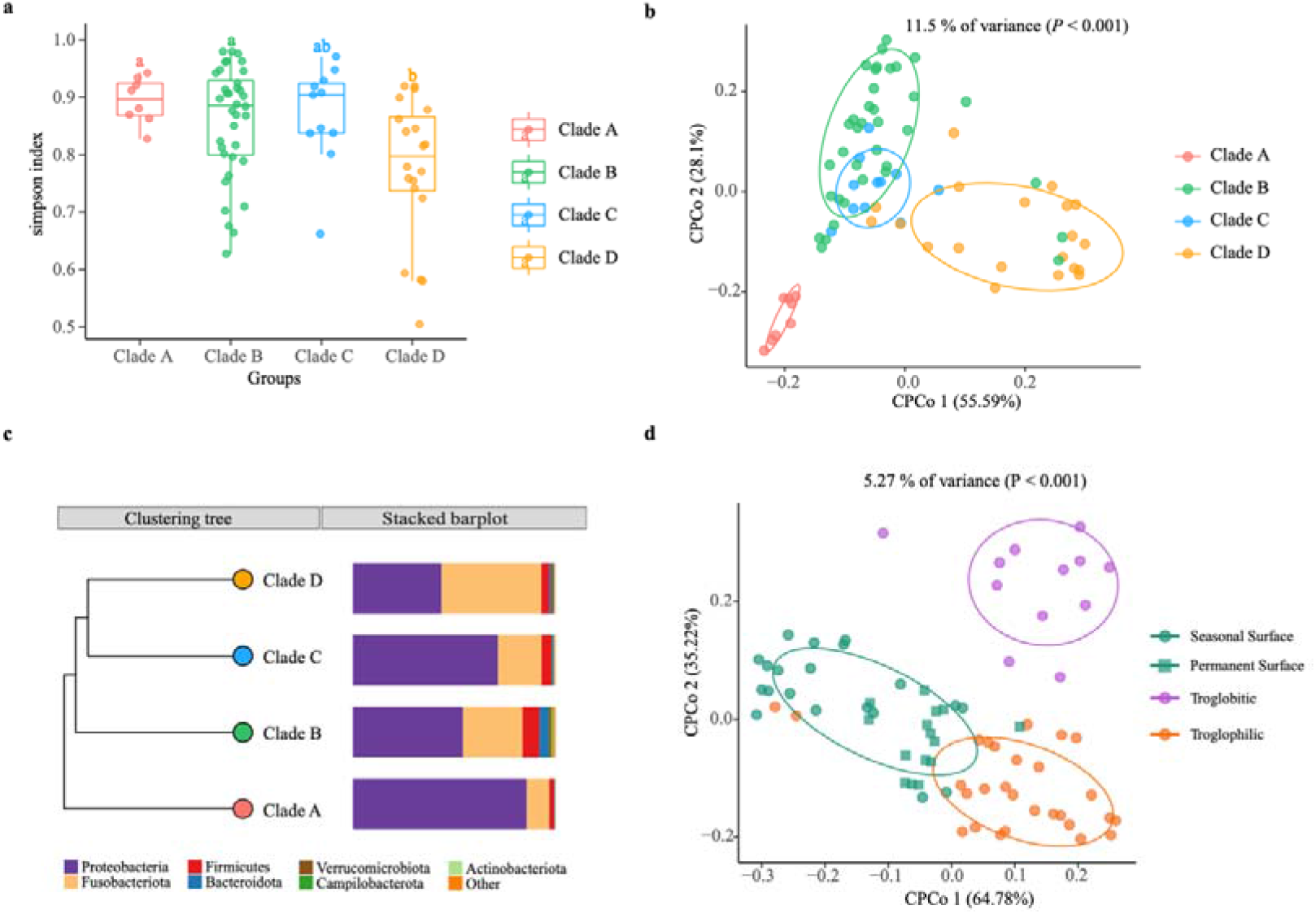
The gut microbiomes of *Sinocyclocheilus* species (N = 24) in context of phylogenetic clades and habitats. **a.** Simpson index. The bottom and top of the box are the first and third quartiles, the band inside the box is the median, and the ends of the whiskers represent the minimum and maximum. Different letters indicate significant difference between groups (ANOVA, Tukey HSD test); **b.** CPCoA of Bray-Curtis dissimilarity showing host phylogeny has significant effect on gut microbiome (11.5% of total variance was explained by the phylogenetic clade, *P* < 0.001). Total number of fishes used: clade A (N = 8), clade B (N = 36), clade C (N = 11) and clade D (N = 20). The ellipses include 68% of samples from each group; **c.** The UPGMA tree with unweighted Unifrac distance matrices representing bacterial community composition of phylogenetic clades at phylum level indicated with bar plots; **d.** CPCoA of Bray-Curtis dissimilarity showing living habitat has significant effect on gut microbiome (5.27% of total variance, explained by the phylogenetic clade, *P* < 0.001). Total number of fishes analyzed: Surface (N = 35), Troglobitic (N = 12), Troglophilic (N = 28). The ellipses include 68% of samples from each group.

### Microbiome community structure associations with host habitat

According to the degree of connection with caves, cavefish living habitats were classified as surface, troglophilic, or troglobitic. We evaluated beta diversity (diversity among samples) using Bray-Curtis distances and did a Canonical Analysis of Principal Coordinates (CAP) to examine the influence of different living habitats on the assembly of *Sinocyclocheilus* cavefish bacterial communities. This analysis revealed that significant differences between samples came from the Surface, Troglobitic and Troglophilic environments, explaining 5.27% of the variation in wild populations (Figure 3d, *P* < 0.001). Troglobitic group is clearly separated from Troglophilic group and the Surface group on the second axis (35.22% variance), while the Surface group is clearly separated from Troglophilic group on the first axis (64.78% variance). CAP analysis showed that the gut microbiota of seasonal surface species (green squares) are more similar to the Troglophilic group compared to Permanent Surface species (green circles). It demonstrates that seasonal Surface species from Clade A and Clade C are similar in gut microbiome structure to Troglophilic species despite their similarity in appearance to permanently Surface species from Yunnan province.

### Influence of surrounding water on the host microbiome

We quantified the contribution of variations in cave water-associated microbiota to the composition of the cavefish gut microbiota. This could signal the effect of a major environmental factor on the assembly of cavefish microbiome.

We analyzed the 19 cave water samples with their corresponding fish samples (Supplementary Table S4). There were 9350 ASVs in the water samples and 4224 ASVs in the fecal samples. The gut microbial ASVs shared with the cave water microbial community accounted for 22.1% (Figure 4a). Next, we used fast expectation-maximization microbial source tracking (FEAST) (Shenhav *et al.*, 2019) to estimate how much of the *Sinocyclocheilus* cavefish gut microbiota derives from cave water. This shows that 52.13% of fish gut microbes came from water sources, while 47.87% came from unknown sources (Figure 4b). Furthermore, the water microbial communities in the 19 caves were also found to be different (Figure 4c). A corollary question is whether the differences in fish gut flora we discovered are attributable to varied microbial exposures in the environment. We assume the causal relationship between these two discrepancies and find two approaches to verify this prediction. First, we focus on whether the cavefish gut microbiota is more similar to the water microbiota of their location, compared to the water microbiota from other locations. The one-tailed t-test, did not show similarity between *Sinocyclocheilus* cavefish gut microbiota and local water flora (Bray-Crutis *t* = −0.094, *P* = 0.5378). Second, we assessed whether the cavefish gut microbiota distance matrix of (18 species, 19 locations) correlated with the distance matrix of the water microbiota between sites. We observed no link between cavefish gut microbiota distance and water microbiota distance using the partial Mantel test, which was adjusted for geographic distance as a confounding variable (Bray-Crutis *R* = 0.066, *P* = 0.275). In conclusion, microbes in cave water serve as a source of gut microbiota for cavefish, but they are not significant enough to explain the differences in gut microbiota among *Sinocyclocheilus* populations in different caves.

**FIGURE 4.**
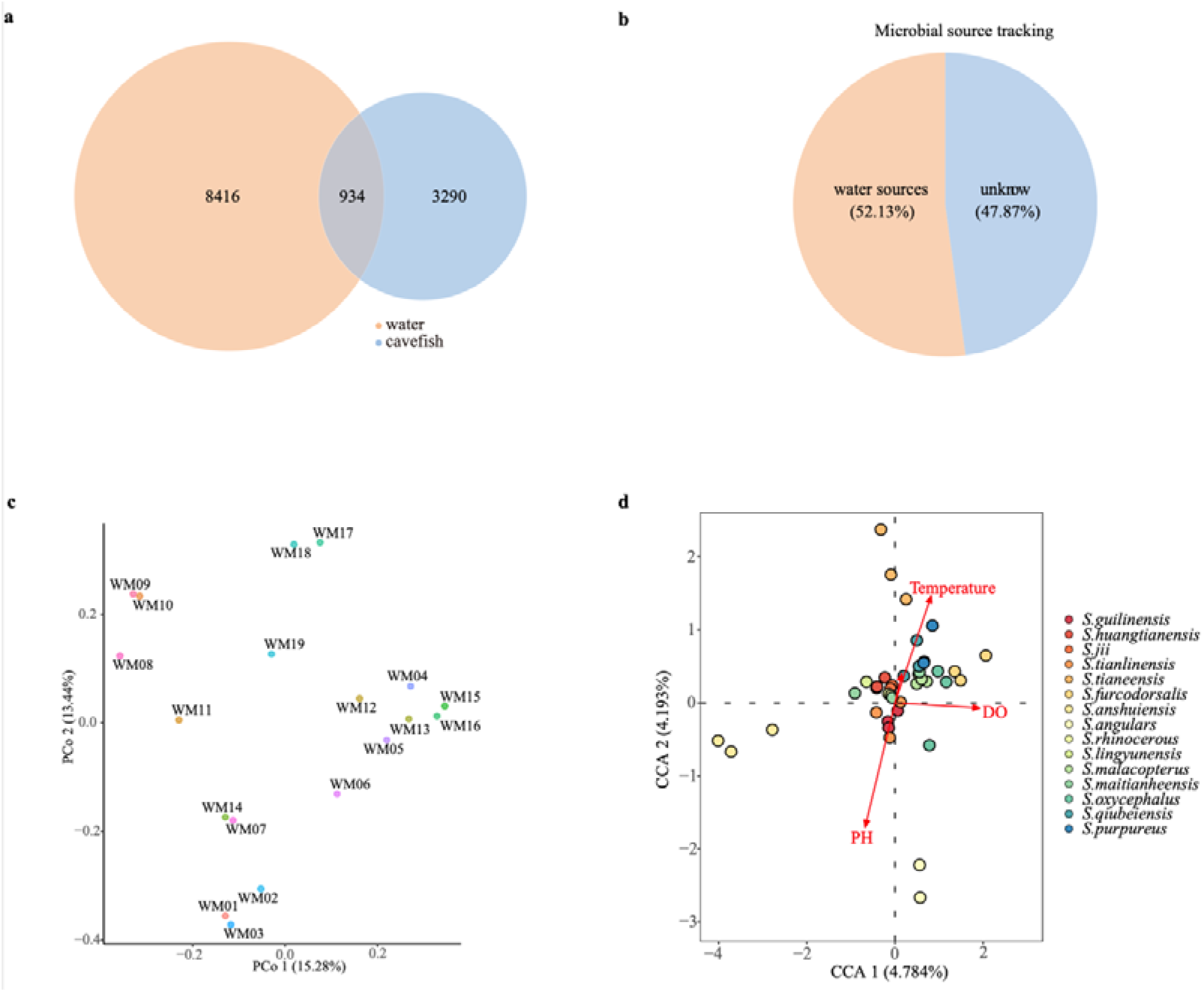
Influence of cave water on the gut microbiome in *Sinocyclocheilus* species. **a.** Number of OTUs shared among cavefishes and water microbiotas.; **b.** Fast expectation-maximization microbial source tracking. Different colors within the circle and area occupied indicate different source in the sink sample; **c.** Unconstrained Principal Coordinate Analysis (Unconstrained PCoA- for principal coordinates PCo1 and PCo2) with Bray-Cutis distance, showing difference between the cave water microbiotas (N = 19); **d.** Water chemistry (changes in the microbial community) explained using Canonical correlation analysis (CCA).

In addition to the microbes in water, water chemistry also may affect the variation of cavefish gut microbiota. We next evaluated the association between physiochemical variables of water and fish gut flora diversity by CCA (Figure 4d). In the CCA analysis, dissolved oxygen (DO), pH and temperature were significantly correlated to cavefish gut microbiota, while conductivity was not. (Figure 4d and Supplementary Table S5). In this analysis, the individual samples show greater separation in the first axis, while DO too, has a higher correlation with the first axis. However, although three physicochemical indicators were related to the shaping of the cavefish gut microbiota, they had a low interpretation rate of 4.3% for DO, 4.2% for pH and 3.2% for Temperature (Supplementary Table S5).

### Changes in the bacterial assemblages from wild to captivity

We collected 24 species of *Sinocyclocheilus* from the wild. Fecal samples were collected from each specimen prior to transfer to captivity. After six months in captivity, fecal samples were again collected from the 17 species that survived over this period. To find changes in bacterial assemblages from wild to captivity, fecal samples from both groups were analyzed (Supplementary Table S6). Measurement of within-sample diversity (alpha diversity) revealed a significant difference between wild and captive populations (*P* <0.001,Tukey HSD test) (Figure 5a and Supplementary Table S7). The fecal microbiota of the captive group, except for three species, had lower richness and diversity than those of the wild group (Figure 5a), indicating that cavefish lost some bacterial species while in captivity. Further, we used non-metric multidimensional scaling (NMDS) analysis with Bray–Curtis distance to investigate the effect of captivity on the intestinal flora of cavefish (Figure 5b and Supplementary Figure S3). The results revealed that the intestinal microbiota of the wild and captive groups divided along the first MDS axis into two distinct clusters: the cluster of captives was more cohesive (Figure 5b), indicating that captivity altered cavefish gut microbiota and, to a certain extent, homogenized the gut bacteria across species.

**FIGURE 5.**
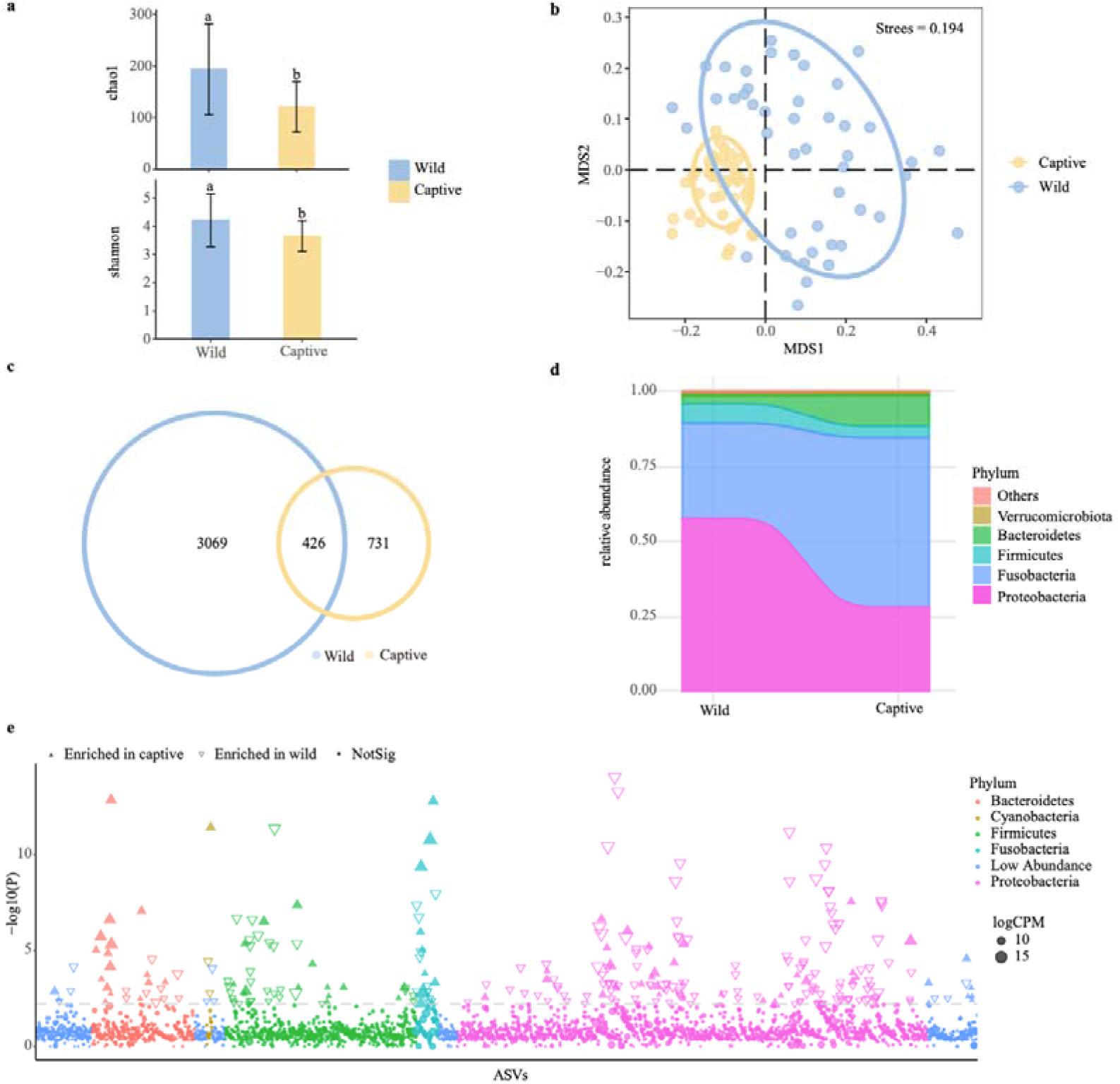
Effect of captivity on the gut microbiota of *Sinocyclocheilus* species (N = 24). **a.** Chao1 and Shannon indices for gut samples from wild and captive populations. Different letters represent significant difference between the groups (*P* < 0.05, ANOVA, Tukey HSD test). Data bars represent mean and error bars represent the standard error of the mean; **b**. Non-metric multidimensional scaling (NMDS) analysis with Bray-Curtis distances, showing gut bacterial composition across the captive and wild cavefishes (*P* < 0.001, ANOSIM test). The ellipses cover 80% of data for each fish sample; **c.** Venn diagram showing unique and shared ASVs between the captive and wild fish populations; **d.** Distribution of gut bacterial taxa in captive and wild fishes at phylum level; **e.** Manhattan plot showing ASVs enriched in the gut of captive and wild fish populations. Each dot or triangle represents a single ASV. ASVs enriched in captive and wild groups are represented by filled or empty triangles, respectively (False discovery rate (FDR) adjusted *P* < 0.05, Wilcoxon rank-sum test). ASVs are arranged in taxonomic order and colored according to the phylum. Counts per million reads mapped (CMP). Replicated samples are as follows: wild group (N = 46), captive group (N = 46). A total of 17 species of *Sinocyclocheilus*.

We discovered not only that captive populations generally had fewer ASVs than wild populations, but also that more than one-third of their ASVs overlapped with wild populations. Hence, in captivity, species of *Sinocyclocheilus* lost a significant portion of their wild-microbiome, but acquired some new ASVs also over the period of six months (Figure 5c and Supplementary Table S7). The total sequences of the captive cavefish group were classified into five major phyla, Proteobacteria, Fusobacteria, Firmicutes, Bacteroidetes, Verrucomicrobiota (Figure 5d). The number of Fusobacteria and Bacteroidetes increased, while the number of Proteobacteria and Firmicutes decreased when compared to the field populations (Figure 5d and Supplementary Table S7). Next, using Manhattan plots, we analyzed the enrichment of ASVs in the wild and in captivity according to their taxonomy (Figure 5e and Supplementary Table S7). ASVs enriched in the captive group belonged to a wide range of bacterial phyla, including Bacteroidetes, Cyanobacteria, Fusobacteria, Spirochaetota and Verrucomicrobiota (FDR adjusted *P* < 0.05, Wilcoxon rank sum test; Figure 5e and Supplementary S7). The wild group had a high abundance of enriched OTUs belonging to Actinobacteriota, Firmicutes and Proteobacteria (FDR adjusted *P* < 0.05, Wilcoxon rank sum test; Figure 5e and Supplementary Table S7). These results suggested that captivity might change the structure of microbial communities in two ways: by removing some ASVs and changing the abundance of shared ASVs. Finally, in order to find fecal microbiota biomarkers that could be used to distinguish wild and captive cavefish, we used the LDA Effect Size (LEfSe) to create a model to test the correlations of wild and captive populations with fecal microbiota data at the phylum, class, order, family, genus, and ASV levels (Supplementary Figure S4).

### Functional prediction of intestinal microbiota and core bacteria in *Sinocyclocheilus*

We used PICRUSt2 software to perform microbiome functional analysis on 75 fecal samples from wild populations and annotated the MetaCyc Metabolic Pathway Database to gain pathway information for 24 species of gut bacteria from provenances (Supplementary Table 8a). We further showed the top 35 pathways in relative abundance using a heatmap plot (Figure 6a). These were involved in three types of biological metabolic pathways: generation of precursor metabolites and energy, biosynthesis and degradation/utilization/assimilation (Figure 6a and Supplementary Table 8b). Surprisingly, gut microbiota in different *Sinocyclocheilus* species appeared to have diverse metabolic pathways, but species from the same phylogenetic clade except for Clade B showed similar pathway abundance (Figure 5a). Among them, Clade A was comparable to Clade C group, whereas Clade D group was distinct from the other clades.

**FIGURE 6.**
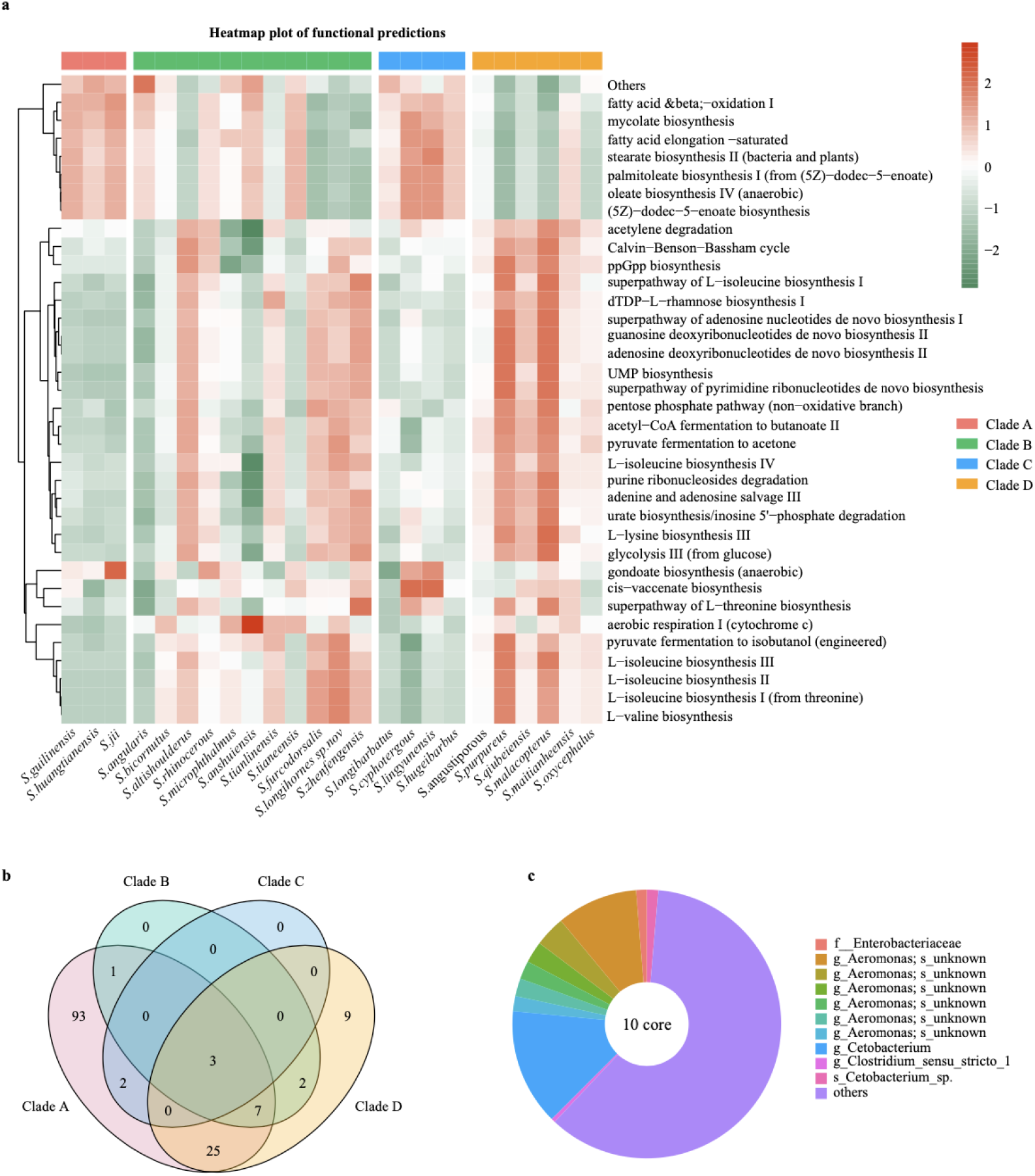
Functional characteristics of differential bacteria and the core microbiome in *Sinocyclocheilus* species (N = 24). a. Heatmap illustrating the metabolic and ecological functions of gut bacteria in different species of cavefishes based on MetaCyc database. The colour scale of higher (red) and lower (blue) shows the functional characteristics in the fish species; b. Venn diagram showing the number of shared and unique OTUs among the *Sinocyclocheilus* species (N = 24) belonging to four clades; c. Diagram representing the core microbiome in *Sinocyclocheilus* species (N = 24).

A follow-up question is whether each phylogenetic clade had a unique microbiota resulting in functional differences. We first identified specific bacteria (Biomaker) important in distinguishing among four phylogenetic clades using LDA Effect Size (LEfSe) method (Supplementary Figure S5). Next, microbiota common to species within each phylogenetic clade are of interest. Species in Clade A shared 131 ASVs, Clade B shared 13 ASVs, Clade C shared 5 ASVs and Clade D shared 46 ASVs (Figures 6b). Combined Figure 6a with Figure 6b, functional prediction of the gut flora showed that the more shared microbiomes the group had the more similar functional prediction of the samples within the group. Note that each phylogenetic clade has an uneven number of species, which may contribute to differences in the number of shared bacteria.

To ascertain the core gut microbes of *Sinocyclocheilus*, we evaluated the shared ASVs across 22 *Sinocyclocheilus* species. Here, we did not use the single samples from *S. cyphotergous* and *S. lingyunensis* as they would have caused us to overlook several microorganisms important to *Sinocyclocheilus* in general. Using the Upset plot, we discovered 10 ASVs across all 22 species., and classifier analysis revealed that 6 ASVs belong to the genus *Aeromonas*, while the other 4 ASVs are f_Enterobacteriaceae, g_Cetobacterium, g_Clostridium_sensu_stricto_1 and s_Cetobacterium_sp. (Supplementary Figure S6 and Table S9a). These shared microbes might constitute the ‘core microbiota’ of the *Sinocyclocheilus* intestine. We next plotted the relative abundance of these 10 ASVs according to their occurrence for the 22 *Sinocyclocheilus* species (Figure 6c and Supplementary Table S9b). We found that although several of these 10 ASVs were discovered in low abundance, in combination, they accounted for around one-third of all gut microbial content (Figure 6c).

## DISCUSSION

We considered the influence of phylogenetic, habitat and environmental factors, including transfer to captivity in shaping the microbiome in *Sinocyclocheilus* cavefishes. From these results, we evaluated the ecological and evolutionary correlates and the adaptability of the microbiome assemblages across the *Sinocyclocheilus* diversification. We hypothesized that the microbiome of *Sinocyclocheilus* is dependent primarily on habitat associations (Troglobitic – Blind; Troglophilic – Micro-Eyed; Surface – Normal) and that phylogeny plays a secondary role. The knowledge from this study is foundational, as a diversification scale analysis of microbiome associations, in which hosts share a MRCA but diverge into multiple novel habitat types, has not been carried out so far.

Despite being a monophyletic and deeply divergent genus of freshwater fishes, the microbiome of *Sinocyclocheilus* showed similarities to the microbiomes of other known fish taxa at a higher taxonomic level. The *Sinocyclocheilus* cavefish gut microbes at the phylum level consisted mainly of Proteobacteria, Fusobacteria and Firmicutes, representing 90% of the community. Others included smaller percentages of Bacteroidetes, Actinobacteria, Verrucomicrobia, Cyanobacteria and Campilobacterota. Our results shows that composition and proportions of the gut microbiota of *Sinocyclocheilus* are similar to those of other Teleost fishes (Llewellyn *et al.*, 2014). Despite the obvious differences in composition among *Sinocyclocheilus* species, the major phyla are still Proteobacteria, Fusobacteria and Firmicutes. This is not unexpected, since the Fusobacteriase phyla have been demonstrated to be the dominant amongst most freshwater and marine fish studied to date (Qian *et al.*, 2021; Xin *et al.*, 2021; Zhang *et al.*, 2021).

### Influence of host phylogeny on microbiome association

It has been shown that host phylogeny influences microbial community structure (Brucker and Bordenstein, 2012). Previous studies have demonstrated the existence of “phylosymbiosis” in some species under laboratory or wild conditions (Brooks *et al.*, 2016; Kohl *et al.*, 2017; Ingala *et al.*, 2018). Yet, in these studies, the habitats were not markedly different from each other. For example, genetic differences between populations of three-spine sticklebacks (*G. aculeatus*) living in different lakes were positively correlated with differences in their gut microbiota (Smith *et al.*, 2015). As predicted, differences in gut microbiota among *Sinocyclocheilus* cavefish species appear to be independent of host genetic differences. We have evaluated several possible explanations for this result: (i) In *Sinocyclocheilus,* even sister species occupy different habitats; hence, it appears that the ecological signature overrides the phylogenetic signal of microbiome diversity. (ii) “phylosymbiosis”, as understood at present, is explained at a higher taxonomic level of the host species, or in different populations of the same species (Phillips *et al.*, 2012). At these two extreme scales, especially under similar habitat conditions among the species considered, it is expected to see a pattern of phylosymbiosis. (iii) Microbes that are in a close association to host phylogeny maybe absent or rarely present in fecal collections. For example, a study comparing fecal and intestinal sampling methods noted that the topology of the host phylogenetic tree and the microbiota similarity clustering tree were identical in the gut contents samples and diverged in the fecal samples (Ingala *et al.*, 2018). Taken as a group, there is only weak support for “phylosymbiosis” among *Sinocyclocheilus* cavefish species, but we found that species within the same phylogenetic clade share a similar microbiome and the phylogenetic relationships between phylogenetic clades were consistent with metagenomic microbial similarity clustering relationships. However, these clades also have a distinct geographic distribution context. Of the four clades, A is at the eastern end of the *Sinocyclocheilus* distribution and D is at the Western end, with B and C overlapping each other in the centre of the distribution. If the microbiomes of clades A and D are very different from each other, and clades B and C are similar, one would expect that distribution of the clade is important for the determination of the microbiome, rather than the phylogeny per se. And this is indeed what we observed. That is, from east to west, Proteobacteria decreases and Fusobacteria increases. Thus, the “phylosymbiosis” exists between phylogenetic clades of *Sinocyclocheilus* cavefish rather than species, and the geographic distribution rather than phylogeny seems to explain abundance of microbial composition.

### Influence of habitat on microbiome association

Habitat occupation is a significant determinant of the microbiome of the cavefishes, and it seems to override the phylogeny and geographic distribution. Previous studies show that habitat type is reflected in eye condition, but with a clear distinction, where one group of Surface fish (Clade D) are permanently surface dwelling (due to lack of caves in the habitats they live in), but in the others (Clade A) where caves are available, they enter caves seasonally when the surface water dries out. *Sinocyclocheilus* have access to more resources as they progress from Troglobitic to Surface (Clade D); hence, the cave environment is thought to have a direct impact on the abundance of food sources available to *Sinocyclocheilus* species. The previous study on the relationship between gut microbial diversity and feeding habits in *Sinocyclocheilus* showed that the gut microbiota of cave dwelling (Troglobitic) species is more conducive to protein digestion and bile secretion, while Surface species has a preference for plant food sources (Chen *et al.*, 2019). Our results demonstrate that there are significant differences in the gut microbiota structure of *Sinocyclocheilus* reflective of the three habitats. As we hypothesized, Troglobitic species have a different gut microbiome structure compared to Troglophilic and surface species. Hence, the unique gut microbiota can be considered as another line of evidence of their adaptation to the cave environment.

### Water as a determinant of *Sinocyclocheilus* gut microbiome

Water may have an impact on the assembly of the fish gut microbial community in two ways: (i) colonization of the fish gut by aquatic microbes (Giatsis *et al.*, 2015) and (ii) the immediate influence of physicochemical parameters of water (Sylvain *et al.*, 2016). Our results suggest that effects from both pathways may exist between cave water and cavefish gut microbiota differences. Because the feeding behaviour of *Sinocyclocheilus* cavefish has not been recorded, our study did not analyse food as a source of microbiota of other potential environmental sources (for example, sediment, bat guano). Surprisingly, our results suggest that about half of the cavefish gut microbiota is formed by colonization with water-associated microbes. A previous study using Bayesian community-level source tracking to estimate the source of gut microbiota in sticklebacks revealed that an average of 12.6% of fish gut microbes were of aquatic origin (Smith *et al.*, 2015), which is markedly lower than our results. The use of fish fecal samples may be one of the reasons for the discrepancy, and it cannot be denied that microbes in fish excreta differ from those in the whole fish intestine (Nielsen *et al.*, 2017). It is worth considering whether fact that more water-derived microbes were present in the feces than water-derived microbes in the gut suggests that some water-derived microbes have not successfully colonized in the fish gut and were excreted (evidence of rejection). Dissolved oxygen (DO), pH and Temperature were demonstrated to influence the assembly of *Sinocyclocheilus* cavefish gut microbes in our data. Coincidentally, DO was also an influential factor in the difference in gut microbiota between the cave population and the surface population of *Astyanax mexicanus* (Ornelas-Garcia *et al.*, 2018). That is, gut microbial community diversity of cavefish appears to be sensitive to water DO concentration.

### Effect of captivity on the fish microbiome

We considered captivity as an extension of the habitat where changes in the microbiome took place as an adaptation to captivity. By comparing wild and captive populations, we found that not only the observed ASV and diversity of the fish gut microbiota were significantly reduced after captivity, but also that the microbiota structure was altered. A plausible explanation is that the fish gut microbiota responded to the environmental change from complex and diverse provenances to a homogeneous environment of the laboratory. We also found that each of the 17 captive species still had a distinct, yet diminished microbiota, despite being in the same recirculating water system and fed on a similar diet. However, the differences in gut microbiota are less evident in laboratory fishes than in wild fishes. For example, NMDS analysis revealed a more similar gut microbial community structure in lab fishes, while in the wild fishes the microbial community structure was relatively dispersed. Since 17 species of *Sinocyclocheilus* cavefishes thrived under laboratory conditions, acquiring a similar microbiome over a long period of time, we conclude that these fishes are adaptable.

A previous study suggested that the effects of captivity on the alpha diversity of fish gut microbes may be species-dependent, such that some species did not change after captivity (Jessica *et al.*, 2016). Our results found a decrease in alpha diversity (Chao1 and Shannon indexes) for most species, but an increase in *S. tianlinensis*, *S. lingyunensis* and *S. maitianheensis*. We observed that there is an approximate alpha diversity index range in captive species: the Chao1 index from 100 to 150 and the Shannon index from 3 to 3.5 (Supplementary Figure S7).

A closer observation of the bacterial phyla also suggests a functional adaptation towards captivity related changes of the microbiome. In captivity, the diversity of Fusobacteria and Bacteroidetes increased, while Proteobacteria and Firmicutes decreased; in fact, Fusobacteria became the most abundant phylum of microorganisms in laboratory fish. Bacteroidetes has been shown to produce digestive enzymes and members of Bacteroidetes have a greater competitive advantage in the gut during periods of relative food deprivation (Xia *et al.*, 2014). Captive fishes were offered food that consisted mainly of protein and fat; this may be responsible for the increase of Bacteroidetes in the intestine. Several studies of the gut microbiota in Cyprinidae have again observed that Fusobacteria is more prevalent in captive fish (Ni *et al.*, 2014; Jessica *et al.*, 2016), so the various microorganisms belonging to Fusobacteria may be more suited to the homogeneous environment of the laboratory. Thus, we support the opinion that not only the fish but also their gut microbiota can be domesticated.

### Core microbiome

In addition to the diverse gut microbiota of *Sinocyclocheilus* cavefish, our faecal bacteria related data from 22 species (nearly one-third of recorded species) point towards 10 ASVs that are shared across *Sinocyclocheilus* cavefish species. Functional roles of these 10 ASVs suggests that they may be important for cavefish to extract essential vitamins and maintain homeostasis. For example, *Cetobacterium* is a symbiotic bacterium that occupies an important ecological niche in the intestinal tract of freshwater fish (Tsuchiya *et al.*, 2008; Ramirez *et al.*, 2018). Members of the genus *Cetobacterium* have been shown to be associated with vitamin B12 synthesis in freshwater fish, as well as in aiding carbohydrate utilization by modulating glucose homeostasis (Wang *et al.*, 2021; Xie *et al.*, 2021). *Aeromonas* is a major colonizer in the gastrointestinal tract in freshwater fish (Trust *et al.*, 1979; Haruo *et al.*, 1985) and these release numerous proteases, thus aiding in digestion (Pemberton *et al.*, 1997). However, some members of the genus *Aeromonas,* such as *A. hydrophila,* are conditionally pathogenic bacteria (Muduli *et al.*, 2021). Although our results did not indicate harmful species of *Aeromonas*, we found a significant reduction of *Aeromonas* abundance in the cavefish gut after captivity.

The bacteria represented by the core microbiome constituting of 10 ASVs did not disappear following captivity, but their compositional abundance increased or decreased. Furthermore, by comparing to a study on the intestinal flora of *Sinocyclocheilus* cavefish, it was found that the bacteria represented by these 10ASVs were also found in the intestines of three species (*S. qujingensis*, *S. aluensis*, *S. lateristritus*) that were not used in our study (Chen *et al.*, 2019). Therefore, a member of family Enterobacteriaceae, six members of the genus *Aeromonas*, two members of the genus *Cetobacterium* and a member of the genus *Clostridium*_sensu_stricto_1 represent a “core microbiota” in *Sinocyclocheilus* cavefish. Given that different species of *Sinocyclocheilus* cavefish contain these glucose homeostasis-regulating and protein-degrading microbes, there may be strong selective pressure to maintain these “core microbes”. Although *Sinocyclocheilus* is monophyletic, its core microbiome does not seem to be specialized at a higher taxonomic level, contrary to what we expected.

*Sinocyclocheilus* species are difficult to sample due to inaccessibility of their habitats, made worse by their small population sizes. Many of the species are known only from their type locality, from a few specimens or drawings from when they were first discovered. Hence an ethical way of sampling adequate numbers of these fishes is through fecal sampling, which we resorted to here. Though fecal sampling does not capture the entire gut microbiome, it allows repeated sampling of the same population. Since *Sinocyclocheilus* fishes are extremely sensitive to stress, we did not attempt abdominal-stroking to force collect faeces as some studies of larger and more robust fishes have done (Jessica *et al.*, 2016).

*Sinocyclocheilus*, allowed us to study microbiome community related adaptations in the largest diversification of freshwater cavefishes in the world. We hypothesized habitat to be the most important determinant of microbiome diversity for this group of fish. Our results show that habitat is indeed important, as is, to a lesser extent, the phylogeny, in determining their gut microbiome. Phylogeny has a geographic context, and we showed that the most divergent clades (A and D) at the Eastern and the Western ends of the *Sinocyclocheilus* distribution respectively show the most divergent microbiomes; the two clades that overlap (B and C) have a similar microbiome. We also show that they acquire a common microbiome in captivity, irrespective of their phylogenetic position, region of origin and habitat, indicating that they are adaptable in the context of microbe related changes in their environment. This further reinforces the importance of habitat as a determinant of the gut microbiome. Finally, the core microbiome of *Sinocyclocheilus* at a higher taxonomic level is remarkably similar to the other teleost fishes, indicating that freshwater fishes maintain a similar microbiome to achieve their physiological needs related to the aquatic realm.

## MATERIALS AND METHODS

### Sample collection, captive care and animal ethics

To address the questions of phylosymbiosis, influence of the environment, captive altered and the core microbiome in this cavefish system, we inventoried the gut microbial communities of 24 species of *Sinocyclocheilus* from provenance to domestication, as well as collecting bacterial samples from cave water. *Sinocyclochelius* species (N = 24) were collected from Guangxi Zhuang Autonomous Region, Guizhou Province and Yunnan Provinces during the period of May, 2020 – December, 2020 (Figure 1). Majority of fishes were collected from caves and few from surface rivers, depending on the species. For each species, live specimens were sampled (N = 3 to 5) using umbrella nets and placed in sterile container and water until a fresh fecal sample was collected (wild-sample). Fecal samples were collected using disposable sterile pipettes and placed in sterile vials. These were immediately frozen and stored in liquid nitrogen at −80 °C until DNA extraction. The collection time for the fecal sample was less than 10 minutes as the excitement after being caught made the fish defecate quickly. Then the fish were placed in polythene bags containing water from the habitat, filled with medical grade oxygen, and transported to laboratory in heat-proof boxes. Due to the rarity of these fishes, note that we used fecal samples as a surrogate for the gut samples to facilitate subsequent captive studies as well.

Water samples from 13 caves were collected from the areas where fishes inhabit. Water samples from seven caves were not collected due to complex (often deep) habitats. One liter of water sample was collected from three sites of a particular area (three liters in total) and stored in sterile bottles. Upon collection, the water samples were filtered through 0.22 μm filter membranes using a portable vacuum pump and immediately frozen and stored in liquid nitrogen at −80 °C until DNA extraction. A portable multi-parameter water quality analyzer (MACH HQ30d) was used to measure conductivity, dissolved oxygen (DO), pH and water temperature, directly from the cave water (ca. 50 cm below the surface).

Fishes collected from field were brought to laboratory and were conditioned for long term captive rearing. They were placed in independent isolation tanks for two weeks before being introduced into the main system. Fish that were injured and whose condition deteriorated over the two weeks were not introduced into the main system. The test subjects whose fecal samples were collected were placed in an ESEN fish rearing system. Each species of *Sinocyclocheilus* were kept in separate box, with a common water circulation system. All laboratory fishes were kept at 18-20°C, were fed ad libitum by a diet based on protein (about 45%) and fat (about 10%). Fecal samples in captivity were collected after six months after being captured. For this, each fish was placed in a sterile plastic box and the fecal sample collected using a sterile dropper as soon as the fishes defecated. Samples collected were again placed in a sterile vial and immediately frozen at −80 °C until DNA extraction.

The present study was conducted in accordance with the recommendations of Institutional Animal Care and Use Committee of Guangxi University (GXU), Nanning-China. All fish were cared for at a fish-room of the Eco.Evo.Devo Group in Forestry college (GXU2019-071).

### DNA extraction, PCR amplification and sequencing

Bacterial genomic DNA was extracted from fecal and water samples according to the protocol set by Novogene Corporation (Beijing, China). Total genomic DNA from samples was extracted using SDS (Sodium Dodecyl Sulfate) method. Concentration of DNA and its purity was monitored on 1% agarose gels. DNA was further diluted to 1 ng/μL using sterile water.

Hyper-variable regions (V3-V4) of the 16S rRNA gene were PCR-amplified from genomic DNA, by using the bacteria-specific universal barcode-primers 341F (5’-CCTAYGGGRBGCASCAG-3’) and 806R (5’-GGACTACNN GGGTATCTAAT-3’). All polymerase chain reactions were performed using 15 μL of Phusion^®^ High-Fidelity PCR Master Mix (New England Biolabs), 0.2 μM of each forward and reverse primer and 10 ng of DNA template. Thermal cycling conditions were as follows: initial denaturation at 98 °C for 1 min, followed by 30 cycles of denaturation at 98 °C for 10 s, annealing at 50 °C for 30 s, elongation at 72 °C for 30 s and final extension at 72 °C for 5 min.

PCR products were mixed in equal volumes of 1X loading buffer containing SYB green and further electrophoresis was operated on 2% agarose gel for detection. PCR products were mixed in equal ratios. The PCR products were further purified using Qiagen Gel Extraction Kits (Qiagen, Germany). Sequencing libraries were generated with NEBNext^®^ Ultra™ IIDNA Library Preparation Kit (Cat No. E7645). Quality of library generated was evaluated on a Qubit@ 2.0 Flurometer (Thermo Scientific) and Agilent Bioanalyzer 2100 system. The library was sequenced on an Illumina NovaSeq platform and 250 bp paired-end reads were generated. Paired-end reads were assigned to samples based on their unique barcodes and were truncated by cutting off the barcodes and primer sequences. Paired-end reads were merged using FLASH (Version 1.2.11) which resulted in raw tags (Magoč *et al.*, 2011). To obtain high quality clean tags, quality filtering on raw tags was performed using fastp software (Version 0.20.0). To detect the chimera sequences, the clean tags were compared with Silva database using Vsearch (Version 2.15.0). The chimera sequences were further removed to obtain effective tags. Raw sequences generated were submitted to the National Center for Biotechnology Information (NCBI) BioProject database (accession number PRJNA772569).

### ASVs denoise and taxonomic annotation

To obtain initial ASVs (Amplicon Sequence Variants) denoise was performed using DADA2 in QIIME2 software (Version 2021.4). ASVs with abundance less than five were filtered out. Taxonomic annotation was performed within Silva Database using QIIME2 software (Version 2021.4). Further, to examine the phylogenetic relationship of individual ASV and the differences of the dominant species among different samples, multiple sequence alignment was performed using QIIME2 software (Version 2021.4). The absolute abundance of ASVs was further normalized using standard sequence number corresponding to the samples with least sequences. Subsequently the alpha and beta diversity analysis were performed based on the output of the normalized data.

### Data analysis and visualization

Alpha diversity indices and beta diversity distances were calculated using QIIME2 software (Version 2021.4). To analyze the diversity, richness and uniformity of communities in each groups, we used box plots to show the Shannon index in R (4.1.1, amplicon package) (Liu *et al.*, 2020). The Tukey’s honestly significant difference test (Tukey’s HSD), Analysis of Variance (ANOVA) test were used for significance testing. To evaluate the effect of different factors on the assembly of bacterial communities, we compared beta diversity using Bray-Curtis distances and performed Canonical Analysis of Principal Coordinates (CAP) in R (4.1.1, amplicon package). To explain the effect of environmental factors on the structure of gut microbial communities Canonical Correspondence Analysis (CCA) was carried out using Canoco software. Venn and Upset diagrams were plotted using TBtools (Chen *et al.*, 2020). Manhattan plot were used to display the ASVs enriched or reduced between the wild group and captive group in R (4.1.1, edgeR and ggplot2 packages). To determine the biomarker, Linear discriminant analysis effect size (LEfSe) was implied using LEfSe software (Version 1.0). The taxa with a log Linear discriminant analysis score (LDA) more than four orders of magnitude were considered. To evaluate the functions of communities in the samples and different groups, PICRUSt2 software (Version 2.4.1) was used for the function annotation analysis.

Extent of phylosymbiosis for these fishes can be tested using the associated between host phylogenies and metagenomic relationships of the gut BM. For this we used data from the most recent phylogeny for *Sinocyclocheilus*. We aligned *cytochrome b* (*cytb*) and *NADH dehydrogenase subuni*t 4 (*ND4*) sequences independently in MUSCLE, implemented in MEGA-X. Then both sequences also are concatenated in MEGA-X. We inferred the maximum likelihood phylogenetic tree with IQTREE software by using ModelFinder with the TIM3+F+I+G4 substitution model and 1,000 rapid bootstrap replicates. We estimated genetic distance matrices for *cytb* and *ND4* genes using Mega X software for 24 species of *Sinocyclocheilus*. Also, some data was derived from a published paper from our laboratory (Mao *et al.*, 2021). Phylogenetic distance matrix was used to perform Mantel tests (R package – vegan 2.5.7) with the host microbial similarity distance matrix. To generate UPGMA cluster trees, Unifrac distance across each microbial sample (Unweighed Pair Group Method with Arithmetic Mean) was used (base R function “hclust”). R package phangorn (Version 2.7.1) was used to test phylogenetic congruence between the host phylogeny and microbiome structure, using Robinson Foulds metric to estimate topological similarity.

## Acknowledgements

We acknowledge the following individuals and organizations: Chenghai Fu, Jiajun Zhou, Yining Chen and Tao Luo for assistance with fish sampling; Hongfu Yang (fisheries workstation in Qubei County, Yunnan Province) for providing us with support and knowledge; Dan Sun and Gajaba Ellepola (Guangxi University) for assistance with data analysis; Rohan Pethiyagoda (Australian Museum) for suggestions to improve the paper.

## Supplementary Information

**FIGURE S1.**
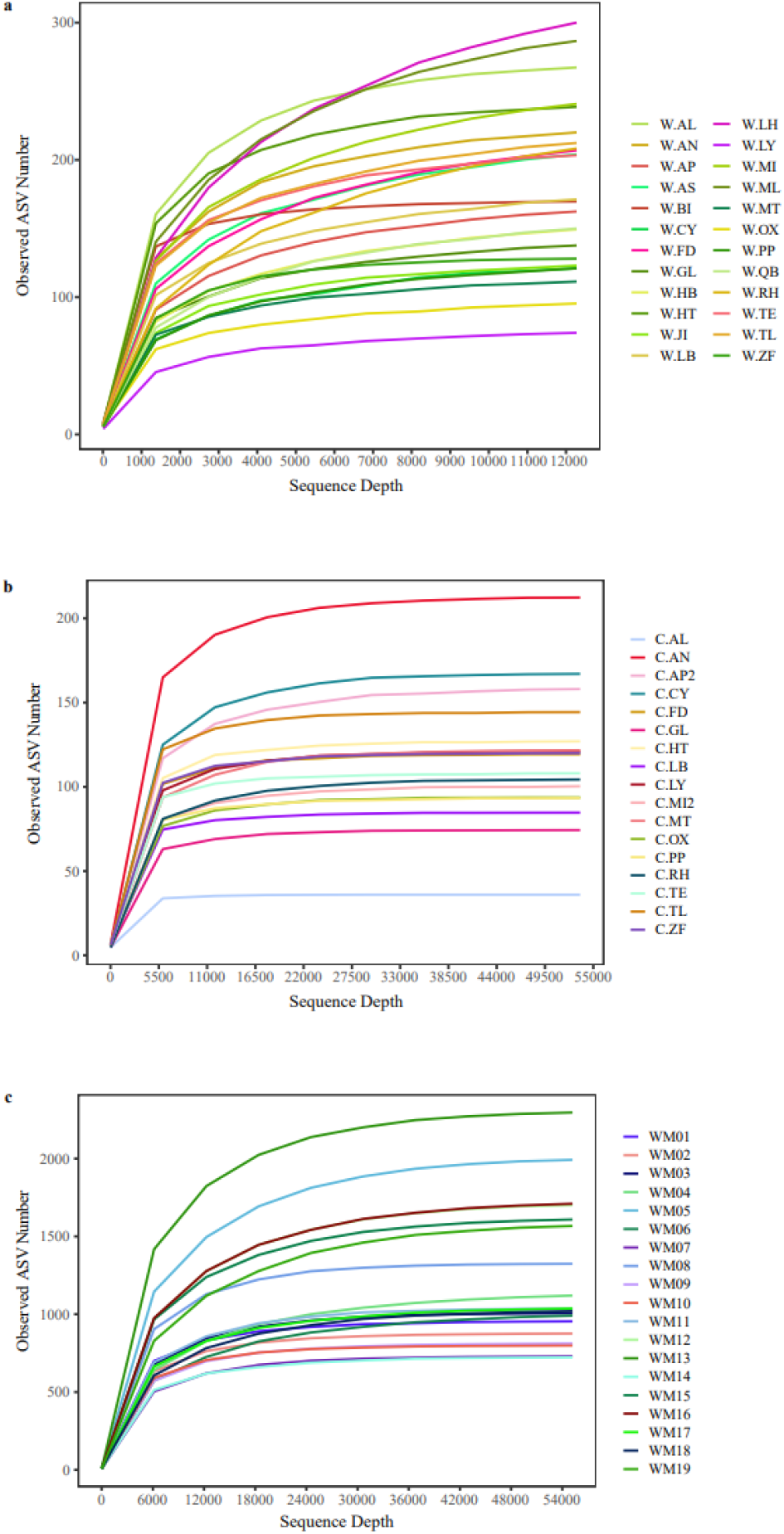
Rarefaction curves of each sample. **a.** Fecal samples from 24 species of wild *Sinocyclocheilus.* **b.** Fecal samples from 17 species of captive *Sinocyclocheilus.* **c.** Water microbial samples from 19 caves. Different colors of the lines indicate different sample, respectively.

**FIGURE S2.**
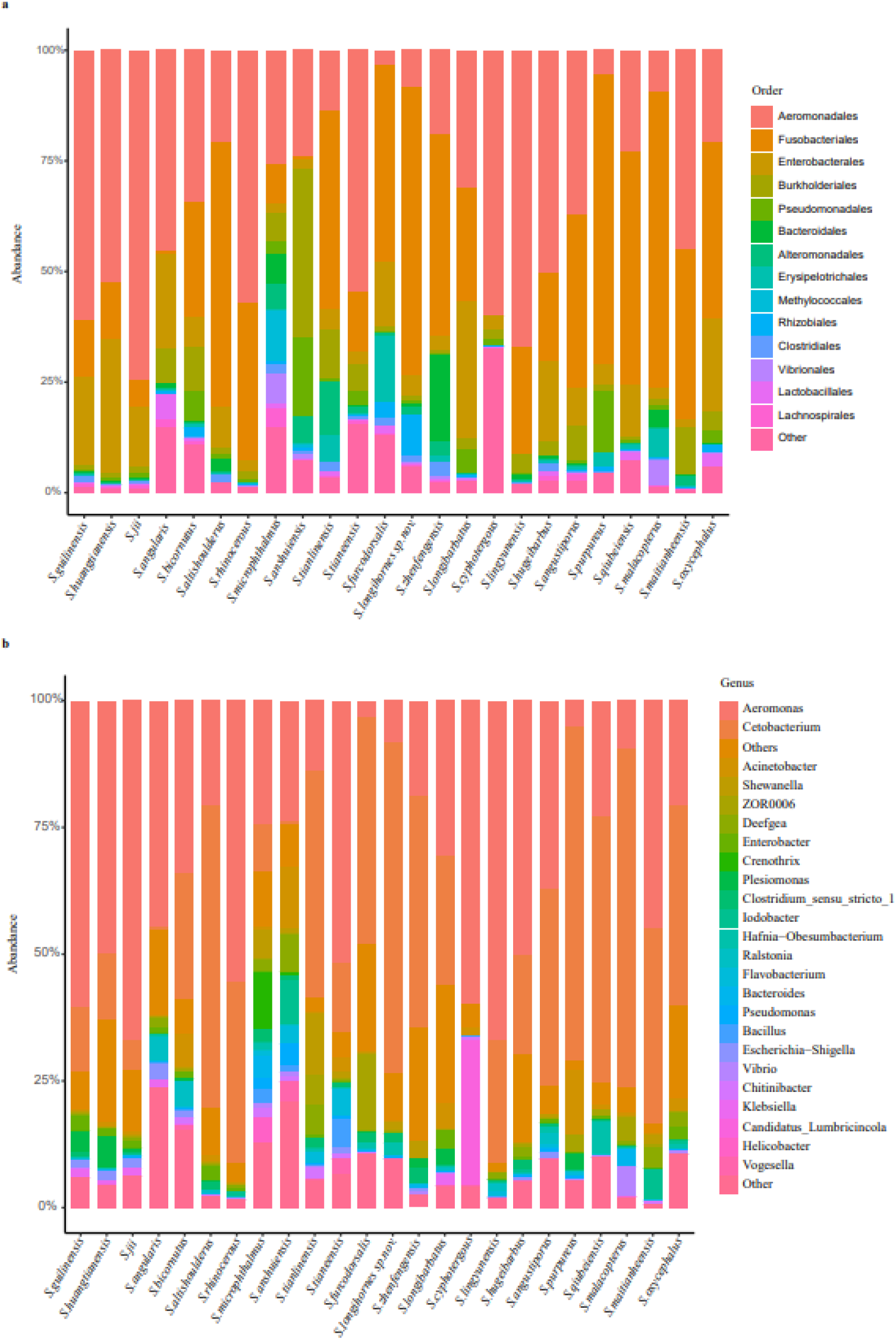
Microbial composition of *Sinocyclocheilus* species. **a.** Order level. **b.** Genus level. Different colors in the figures indicate the different groups, and details are shown on the right sides of each figure, respectively.

**FIGURE S3.**
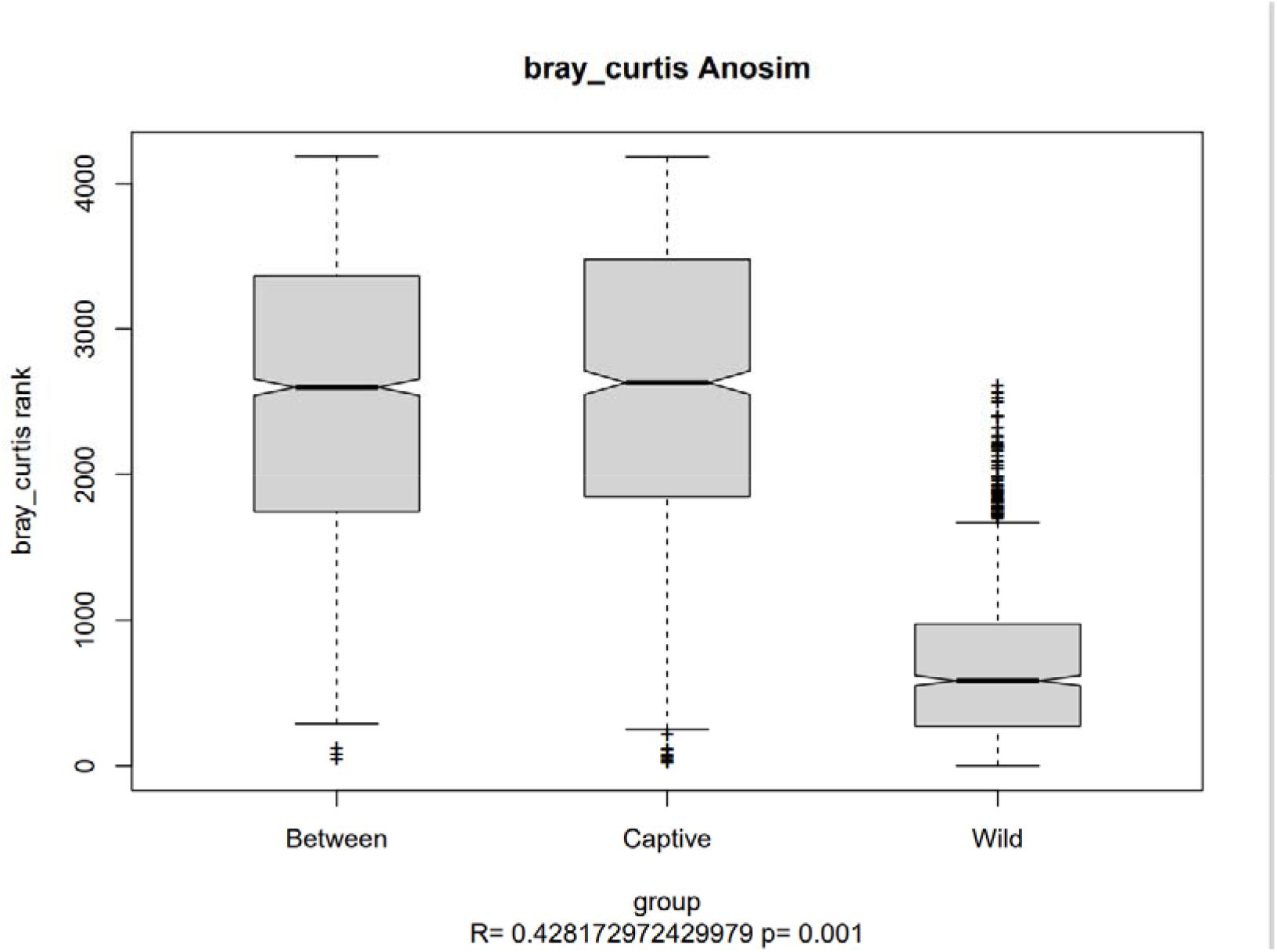
Analysis of similarities (Anosim) test based on Bray-Curtis distance for wild group and captive group.

**FIGURE S4.**
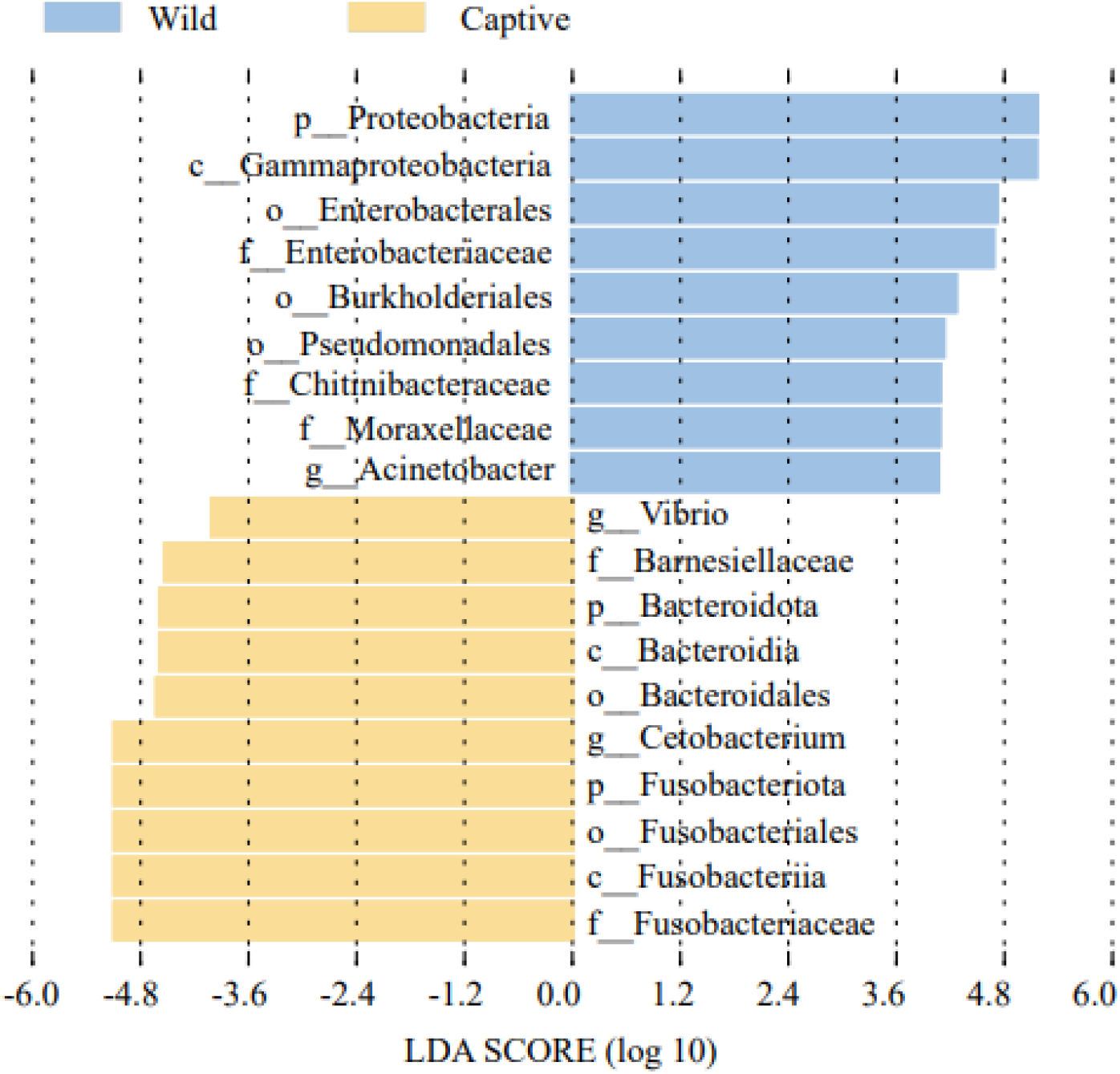
LDA Effect Size (LEfSe) method to find out microbiota biomarkers in wild and captive cavefish.

**FIGURE S5.**
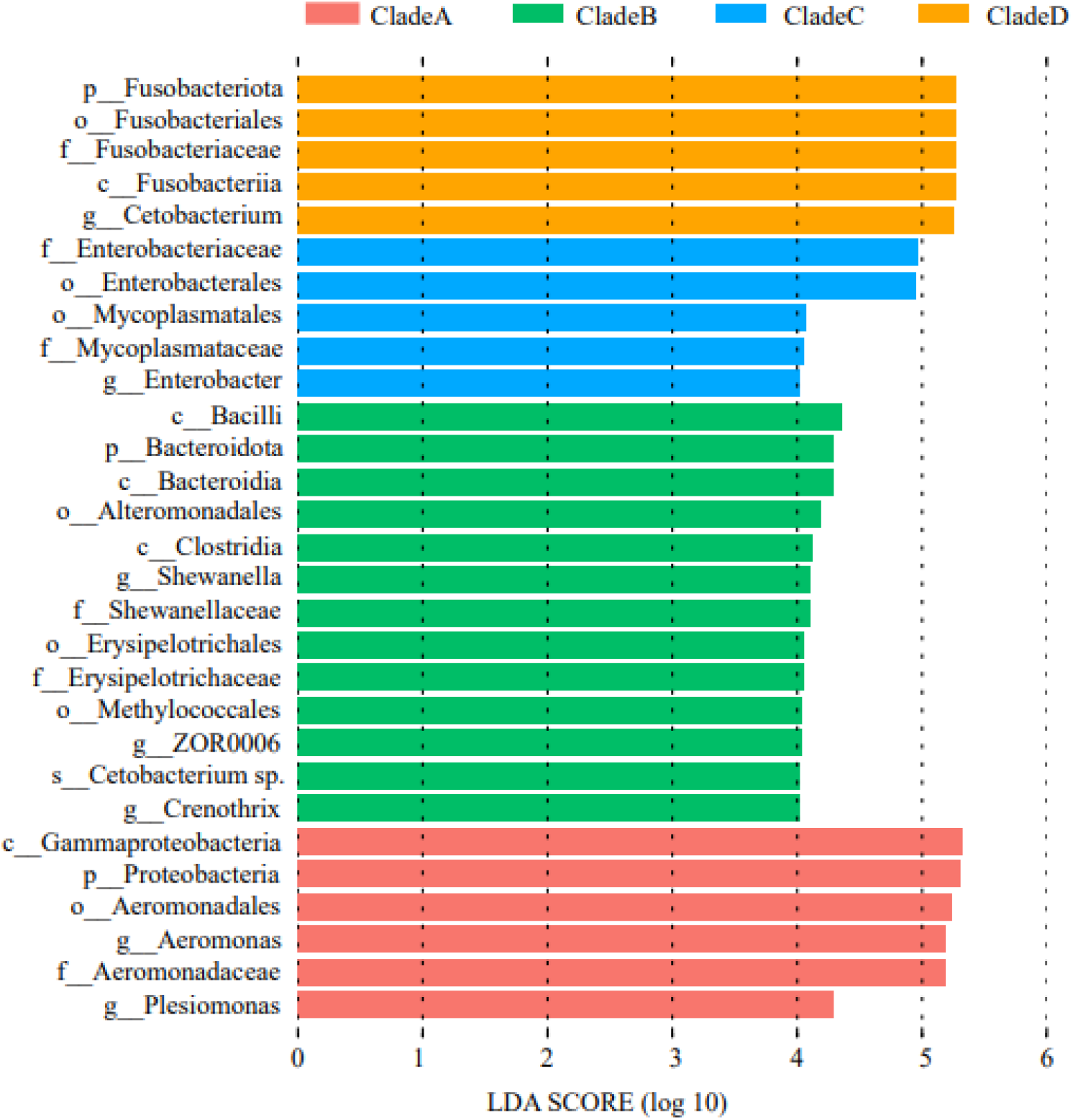
LDA Effect Size (LEfSe) method to find out microbiota biomarkers between four Phylogenetic clades.

**FIGURE S6.**
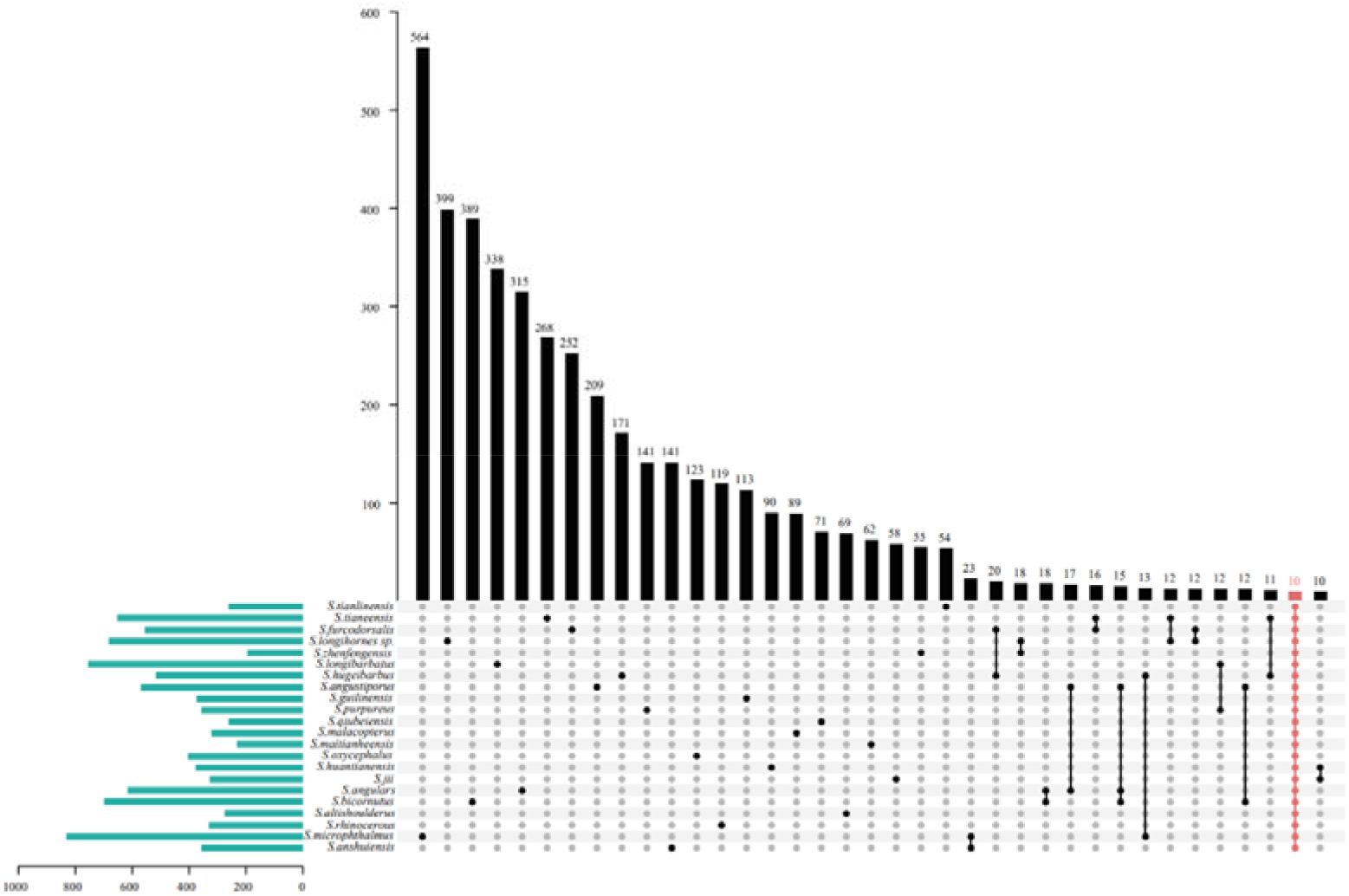
10 shared ASVs among 22 species of *Sinocyclocheilus.*

**FIGURE S7.**
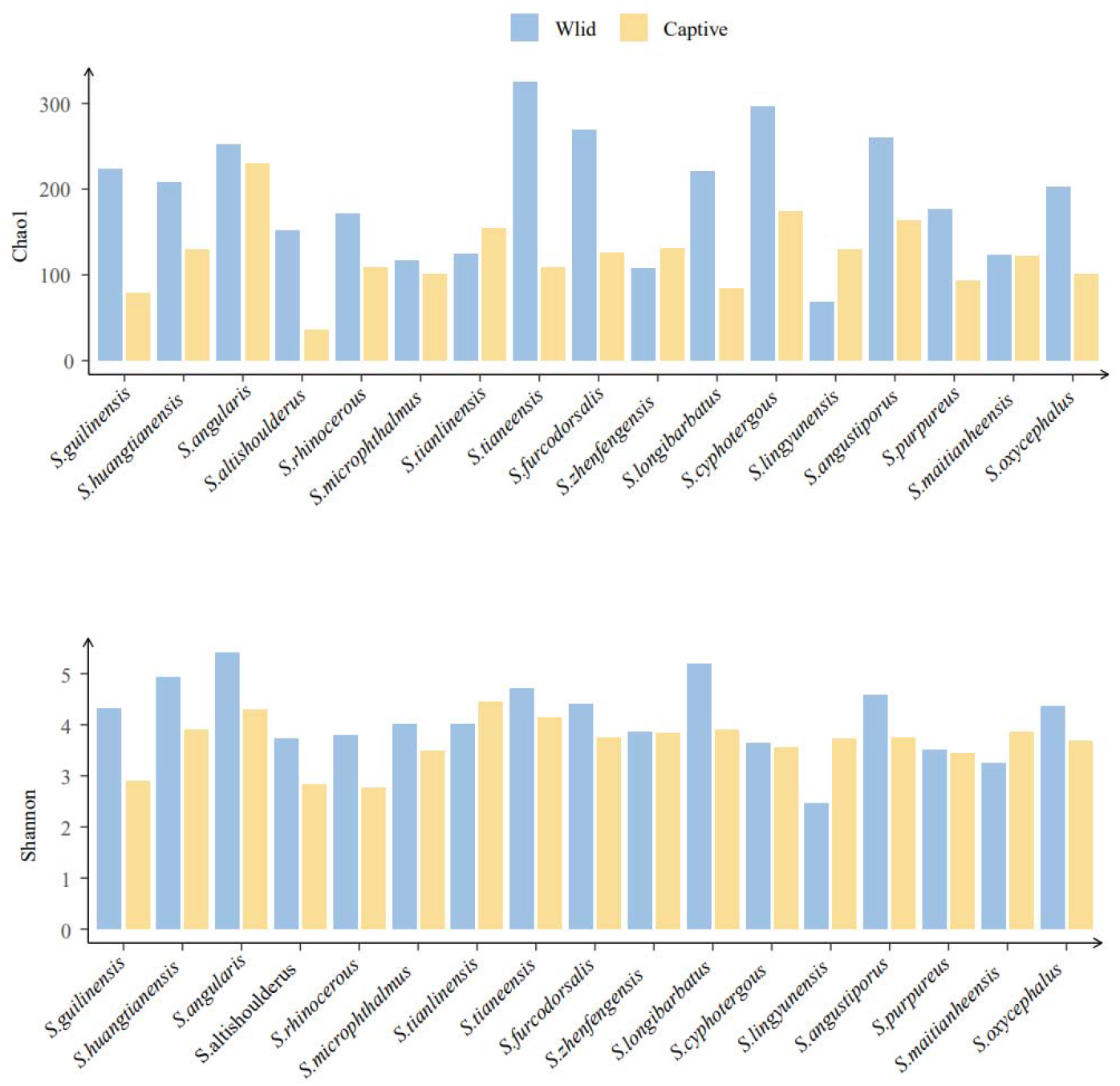
Comparison of alpha diversity indices of *Sinocyclocheilus* (N=17) before and after captivity.

**Supplementary Table S1.**
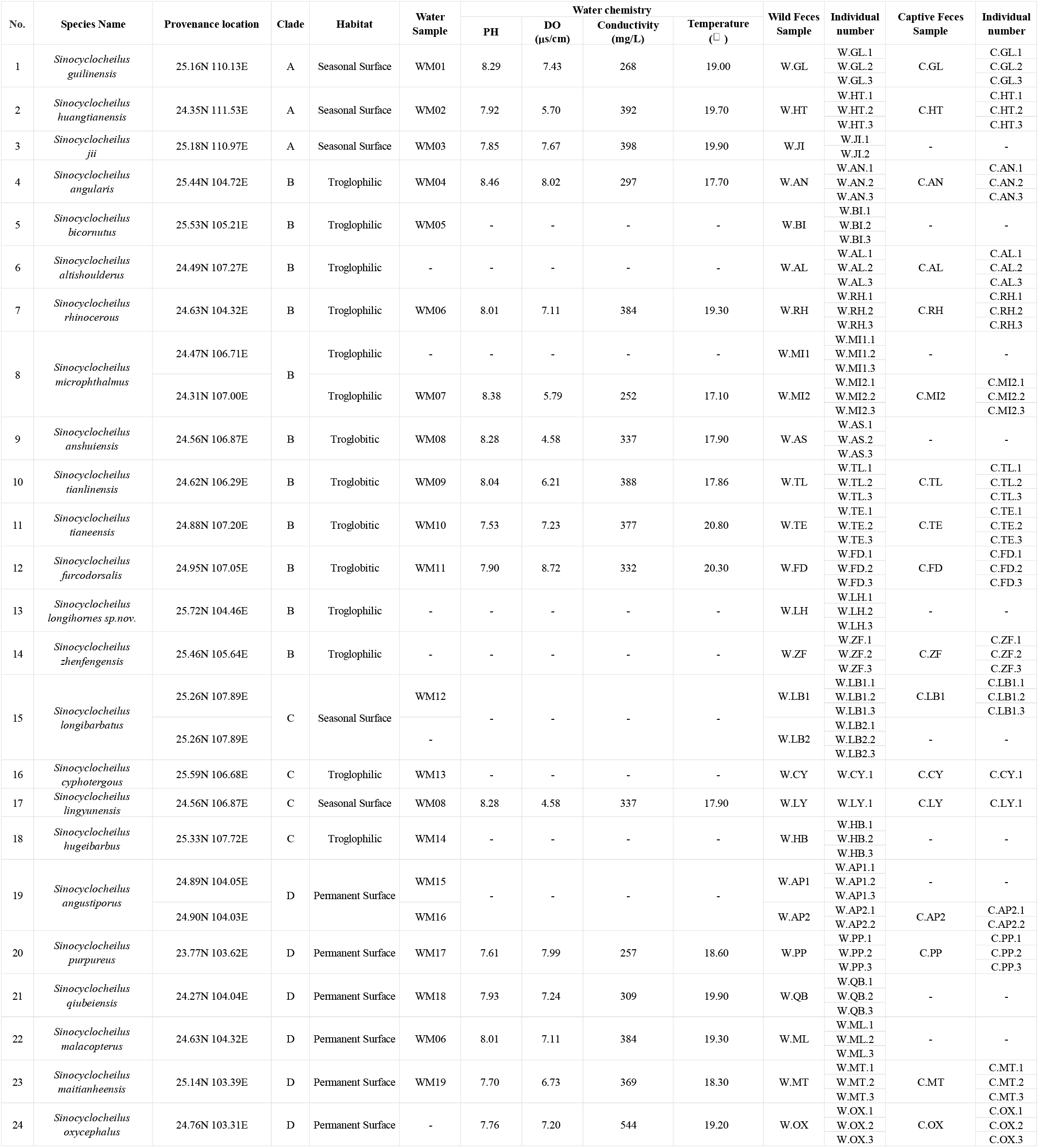
Sample Collection Information Form.

## These Tables are not provided here due to size, but will be provided upon decision to review

Supplementary Table S2 ASV table, Taxonomic annotation and metadata table of wild samples.

Supplementary Table S3 ANOVA, Tukey HSD test of phylogenetic clades.

Supplementary Table S4 ASV table, Taxonomic annotation and metadata table of water and fish microbiota.

Supplementary Table S5 Water chemistry with the results of its analysis.

Supplementary Table S6 ASV table, Taxonomic annotation and metadata table of Captive population.

Supplementary Table S7 Changes in the bacterial assemblages from wild to captivity. 7a. ANOVA, Tukey HSD test of wild and captive groups; 7b. Overlapped 426 ASVs in Wild and Captive populations; 7c. Relative abundance of the two groups at the Phylum level; 7d. Differential abundance of ASVs between Wild and Captive populations.

Supplementary Table S8 Pathway information annotated the MetaCyc Metabolic Pathway Database.

Supplementary Table S9 10 core microbiome of *Sinocyclocheilus* and their abundance.

